# Hypoxia-tolerant vascular endothelial growth factor-A (VEGF-A) and the genetic variations of *VEGFA* in Sherpa highlanders

**DOI:** 10.1101/2020.03.16.993568

**Authors:** Yunden Droma, Takumi Kinjo, Shuhei Nozawa, Nobumitsu Kobayashi, Masanori Yasuo, Yoshiaki Kitaguchi, Masao Ota, Buddha Basnyat, Masayuki Hanaoka

**Author notes:** Corresponding author (MH), (YD). These authors contributed equally to this work (Yunden Droma, Masayuki Hanaoka).

## Abstract

Sherpa highlanders demonstrate extraordinary tolerance to hypoxia at high altitudes, partly by one of the adaptation mechanisms promoting increases of microcirculatory blood flow and capillary density at high altitude for restoring oxygen supply to tissues. Hypoxia stimulates vascular endothelial growth factor (VEGF), which is an important signaling protein involved in hypoxia-stimulated vasculogenesis and angiogenesis. Our present study included 51 Sherpas dwelling in Namche Bazaar village (3440 m) and 76 non-Sherpa lowlanders residing in Kathmandu (1300 m) in Nepal. In these participants, we measured plasma VEGF-A concentrations and genotyped five single-nucleotide polymorphisms (SNPs) of *VEGFA*: rs699947, rs8333061, rs1570360, and rs2010963 in the 5′-untranslated region (5′-UTR); and rs3025039 in the 3′-UTR. The average circulating VEGF-A level in Sherpas did not respond to hypoxia at the high altitude in 3440 m, remaining equivalent to the level in non-Sherpa lowlanders at low altitude. Allele discriminations for the analyzed SNPs revealed significant genetic divergences of rs699947, rs8333061, and rs2010963 in Sherpa highlanders compared with non-Sherpa lowlanders, East Asians, South Asians, and the global population; however, consistency with the indigenous Tibetan highlanders from the Tibet Plateau. On the other hand, the SNP rs3025039 in the 3′-UTR presented constant preserved genetic variation among global populations. Our findings indicated that the physiological sea-level VEGF-A concentration in Sherpa highlanders at high altitude was probably linked with the significant variations of *VEGFA* in Sherpas that regulate the gene expression in a manner of tolerance to hypoxia through production of the optimal biological level of VEGF-A at high altitudes. Precise angiogenesis at high altitude contributes to the adaptive levels of capillary density and microcirculation, providing efficient and effective diffusion of oxygen to tissues and representing human adaptation to high-altitude hypoxia environment.

**Author summary:** Sherpa highlanders demonstrate extraordinary tolerance to hypoxia at high altitudes, partly by one of the adaptation mechanisms promoting increases of microcirculatory blood flow and capillary density at high altitude for restoring oxygen supply to tissues. Vascular endothelial growth factor (VEGF) is mainly stimulated by hypoxia, and is an important signaling protein involved in hypoxia-stimulated vasculogenesis and angiogenesis. Interestingly, we found that the circulating VEGF-A level in Sherpa highlanders did not respond to hypoxia at high altitude. Furthermore, allele discrimination of the single nucleotide polymorphisms (SNPs) of *VEGFA* revealed significant divergences of rs699947, rs8333061, and rs2010963 within the *VEGFA* regulation region in Sherpa highlanders compared to the non-Sherpa lowlanders, East Asians, South Asians, and the global population; however, consistency with Tibetan highlanders from the Tibet Plateau. We propose that the hypoxia-tolerant circulating VEGF-A level in Sherpa highlanders is linked with the genetic variations of *VEGFA*, contributing to human adaptation to high-altitude hypoxic environments.

## Introduction

At altitudes of >2,500 m above sea level, the atmospheric oxygen tension is reduced by >70% compared to at sea level, which is a life-threatening environment for humans originally from lowlands [1, 2]. However, over 140 million people worldwide permanently reside at high altitudes without health issues related to oxygen insufficiency [3]. For example, the Sherpas are a minority ethnic population that have permanently lived along the Himalayan region in Asia for hundreds of generations. Physiologically, the Sherpa exhibit extraordinary tolerance to hypoxia at high altitudes, which is highly valuable during expeditions to extreme altitudes in the Himalayan region [4]. It has been widely hypothesized that genetic adaptation to high-altitude hypoxia, as a consequence of natural selection, may explain their extraordinary physical performances at high altitudes [5, 6].

Hypoxia is generally an important regulator of blood vessel tone and structure, and a potent stimulus for vasculogenesis (*de novo* formation of the embryonic circulatory system) and angiogenesis (growth of blood vessels from pre-existing vasculature) [7]. Hypoxia-stimulated physiological vasculogenesis and angiogenesis may increase microcirculatory blood flow and capillary density, thus restoring the oxygen supply in tissues to compensate for oxygen insufficiency [8]. Compared to non-Sherpa lowlanders, Sherpa highlanders show significantly greater capillary density in muscle cross-section [9], and higher sublingual microcirculatory blood flow and greater capillary density at high altitude [10]. These findings indicate efficient and effective oxygen transport to tissues in Sherpa highlanders, supporting the notion that microcirculation plays a critical role in the mechanism underlying adaptation to hypoxia [11].

Vascular endothelial growth factor-A (VEGF-A) is one of the members in VEGF family and is an endothelial cell-specific angiogenic inducer that has been implicated in multiple processes, including angiogenesis, vasculogenesis, and vascular endothelial cell division [12]. It is a key mediator of downstream production in the hypoxia-response biological pathway, and upregulates vascular smooth muscle and endothelial cells under hypoxic conditions *in vivo* and *in vitro* [13]. Studies of high-altitude medicine have revealed significantly increased circulating VEGF-A levels in patients with acute mountain illnesses, including acute mountain sickness (AMS) [14], high-altitude pulmonary edema (HAPE) [15], and high-altitude cerebral edema (HACE) [16]. Moreover, VEGF-A level is negatively correlated with the decreased percutaneous arterial oxygen saturation (SpO_2_) in individuals susceptible to chronic mountain sickness (CMS) at high altitudes [17]. CMS is a syndrome representing failure to adapt to high altitude—characterized by hypoxemia, blood vessel proliferation, excessive polycythemia, and pulmonary hypertension—which mainly occurs in immigrant lowlanders and occasionally in native highlanders at high altitudes [3]. The unusually increased VEGF-A production at high altitudes pathophysiologically contributes to the formation of abnormal new blood vessels, pulmonary vessel remodeling, vascular smooth muscle cell proliferation [17, 18], and distal vasculogenesis in skin and mucosa [19]— each of which is a potential contributor to CMS development [3].

We hypothesized that VEGF-A may play a role in the mechanism of adaptation to high-altitude hypoxia among Sherpa highlanders. Our results supported that the proper response to hypoxia of VEGF-A in Sherpa highlanders was highly linked with the significant genetic variations of the VEGF-A gene (*VEGFA*), likely as a consequence of natural selection throughout hundreds of generations of exposure to the high-altitude environment.

## Results

### Phenotypes of Sherpa highlanders at high altitude and non-Sherpa lowlanders at low altitude

The Sherpa highlander group comprised 51 Sherpas living in Namche Bazaar village (3440 m) in the Khumbu region, and the Non-Sherpa lowlander group comprised 76 non-Sherpa lowlanders living in Kathmandu (1300 m) in Nepal. The two groups were matched in terms of sex ratio and average age (P > 0.05, Table 1). The average SpO_2_ level was 96.7 ± 0.2% (median, 97%; range, 92–100%) in non-Sherpas at 1330 m, and was 93.7 ± 0.2% (median, 94%; range, 90–97%) in Sherpas at 3440 m. The average SpO_2_ level significantly differed between the two groups due to the different altitudes (P = 6.43 × 10^−17^). However, these groups exhibited similar average plasma VEGF-A levels: 262.8 ± 17.9 pg/m in Sherpas at 3440 m vs. 266.8 ± 21.8 pg/m in non-Sherpas at 1330 m (P = 0.88). The plasma VEGF-A concentration was not significantly correlated with SpO_2_ level in Sherpas at high altitude (r = 0.13, P > 0.05, Fig 1A), but showed a moderate negative correlation with SpO_2_ level in the non-Sherpas at low altitude (r = −0.27, P < 0.05, Fig 1B).

**Table 1.** Phenotypes of Sherpas at high altitude (3440 m) and non-Sherpas at low altitude (1300 m)

| Phenotypes | Sherpa<br>highlanders at<br>3440 m | Non-Sherpa<br>lowlanders at<br>1300 m | P* |
| --- | --- | --- | --- |
| No. subjects, n | 51 | 76 |  |
| Males: Females, n | 21:30 | 40:36 | 0.27 |
| Age, years | 30.9 ± 1.1 | 29.9 ± 0.8 | 0.251 |
| Oxygen saturation, % |  |  |  |
| Mean | 93.7 ± 0.2 | 96.7 ± 0.2 | $6.43 \times 10^{-17}$ |
| Median | 94 | 97 |  |
| Minimum–Maximum | 90–97 | 92–100 |  |
| VEGF-A (pg/mL) |  |  |  |
| Mean | 262.8 ± 17.9 | 266.8 ± 21.8 | 0.88 |
| Median | 247.1 | 202.9 |  |
| Minimum–Maximum | 91.2–644.2 | 59.4–872.8 |  |

Continuous data are expressed as mean ± standard of error (SE).
\* Differences between the two groups were analyzed by Student's t-test for continuous data and by contingency table ( $2 \times 2$ ) for categorical data.

**Fig 1.**
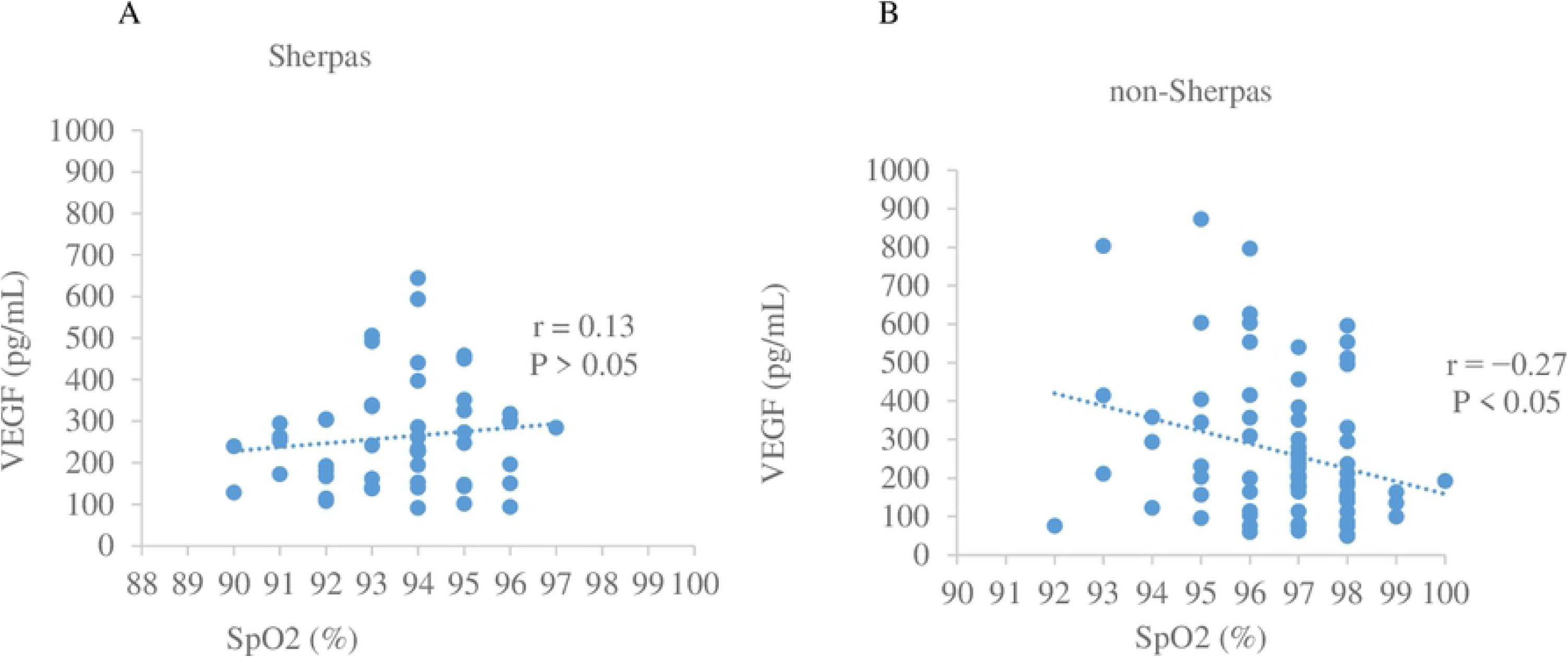
Correlation of plasma VEGF-A concentration with oxygen saturation (SpO_2_) in the two study populations. A) Plasma VEGF-A concentration was not significantly correlated with SpO_2_ level in the Sherpas at high altitude (r = 0.13, P > 0.05). B) Plasma VEGF-A concentration was negatively correlated with SpO_2_ level in the non-Sherpas at low altitude (r = −0.27, P < 0.05).

### Distributions of the SNPs in Sherpa highlanders versus non-Sherpa lowlanders

The genotype distributions and allele frequencies of the SNPs were all met the Hardy-Weinberg equilibrium (HWE) in both groups. Two single-nucleotide polymorphisms (SNPs) in the promoter region (rs699947 and rs833061) and one SNP in the 5′-untranslated region (5′-UTR) (rs2010963) exhibited significant between-group differences in terms of genotype distribution (Pc = 1.43 × 10^−3^, 2.48 × 10^−3^, and 1.04 × 10^−7^, respectively) and allele frequency (Pc = 3.30 × 10^−5^, 4.95 × 10^−4^, and 1.19 × 10^−7^, respectively) (Table 2). On the other hand, the SNP in the 3′-UTR (rs3025039) did not significantly differ between groups with regards to genotype distribution (Pc = 3.65) or allele frequency (Pc = 2.15) (Table 2). The major alleles of these SNPs in the Sherpa highlanders were rs699947**C**, rs833061**T**, rs1570360**G**, rs2010963**C**, and rs3025039**C**.

**Table 2.**
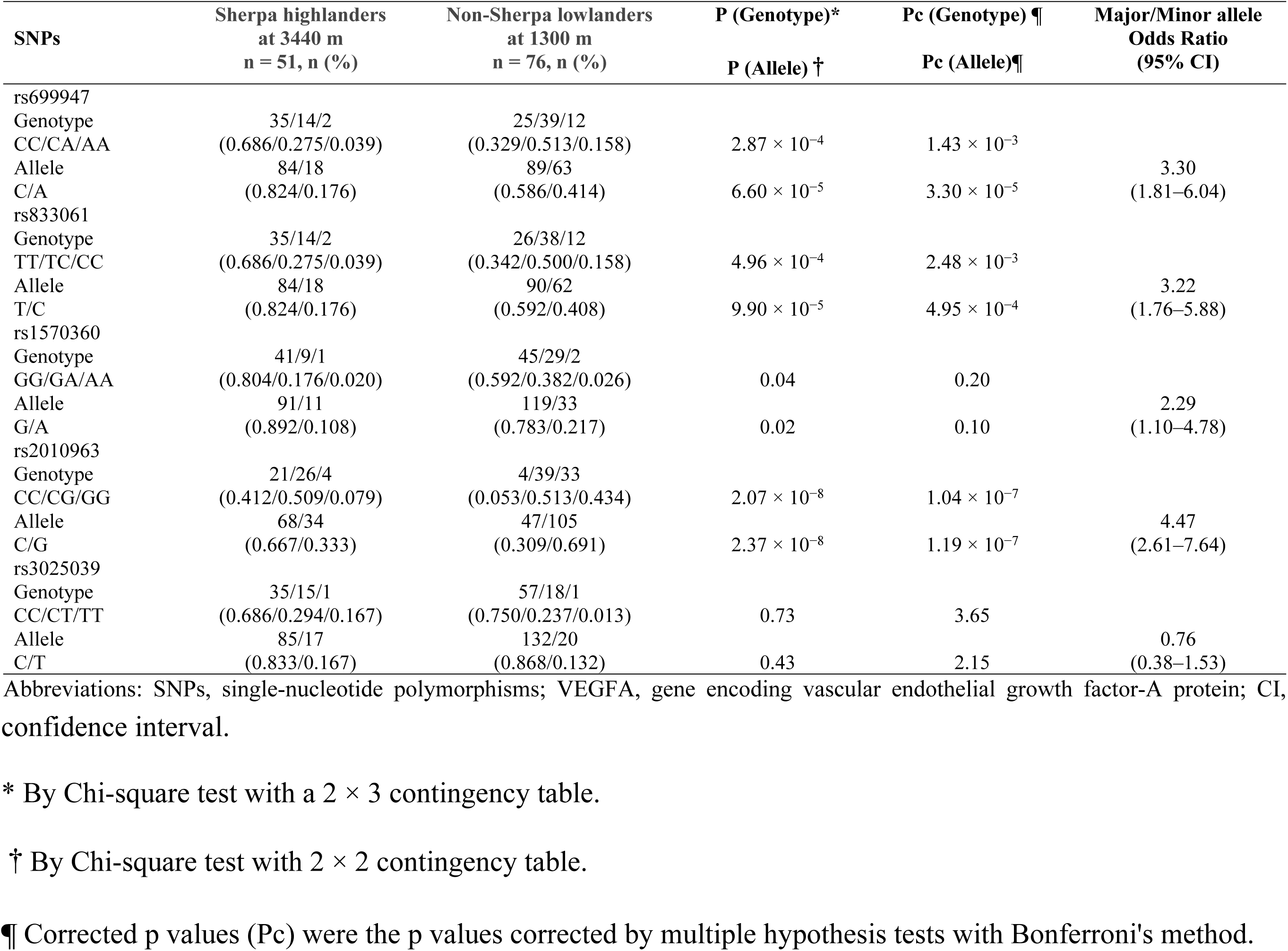
Genotype distributions and allele frequencies of the analyzed SNPs in *VEGFA* among Sherpa highlanders versus non-Sherpa lowlanders.

| SNPs | Sherpa highlanders<br>at 3440 m<br>n = 51, n (%) | Non-Sherpa lowlanders<br>at 1300 m<br>n = 76, n (%) | P (Genotype)*<br>P (Allele) † | Pc (Genotype) ¶<br>Pc (Allele)¶ | Major/Minor allele<br>Odds Ratio<br>(95% CI) |
| --- | --- | --- | --- | --- | --- |
| rs699947 |  |  |  |  |  |
| Genotype | 35/14/2 | 25/39/12 |  |  |  |
| CC/CA/AA | (0.686/0.275/0.039) | (0.329/0.513/0.158) | $2.87 \times 10^{-4}$ | $1.43 \times 10^{-3}$ | |
| Allele | 84/18 | 89/63 |  |  | 3.30 |
| C/A | (0.824/0.176) | (0.586/0.414) | $6.60 \times 10^{-5}$ | $3.30 \times 10^{-5}$ | (1.81–6.04) |
| rs833061 |  |  |  |  |  |
| Genotype | 35/14/2 | 26/38/12 |  |  |  |
| TT/TC/CC | (0.686/0.275/0.039) | (0.342/0.500/0.158) | $4.96 \times 10^{-4}$ | $2.48 \times 10^{-3}$ | |
| Allele | 84/18 | 90/62 |  |  | 3.22 |
| T/C | (0.824/0.176) | (0.592/0.408) | $9.90 \times 10^{-5}$ | $4.95 \times 10^{-4}$ | (1.76–5.88) |
| rs1570360 |  |  |  |  |  |
| Genotype | 41/9/1 | 45/29/2 |  |  |  |
| GG/GA/AA | (0.804/0.176/0.020) | (0.592/0.382/0.026) | 0.04 | 0.20 |  |
| Allele | 91/11 | 119/33 |  |  | 2.29 |
| G/A | (0.892/0.108) | (0.783/0.217) | 0.02 | 0.10 | (1.10–4.78) |
| rs2010963 |  |  |  |  |  |
| Genotype | 21/26/4 | 4/39/33 |  |  |  |
| CC/CG/GG | (0.412/0.509/0.079) | (0.053/0.513/0.434) | $2.07 \times 10^{-8}$ | $1.04 \times 10^{-7}$ | |
| Allele | 68/34 | 47/105 |  |  | 4.47 |
| C/G | (0.667/0.333) | (0.309/0.691) | $2.37 \times 10^{-8}$ | $1.19 \times 10^{-7}$ | (2.61–7.64) |
| rs3025039 |  |  |  |  |  |
| Genotype | 35/15/1 | 57/18/1 |  |  |  |
| CC/CT/TT | (0.686/0.294/0.167) | (0.750/0.237/0.013) | 0.73 | 3.65 |  |
| Allele | 85/17 | 132/20 |  |  | 0.76 |
| C/T | (0.833/0.167) | (0.868/0.132) | 0.43 | 2.15 | (0.38–1.53) |

Abbreviations: SNPs, single-nucleotide polymorphisms; VEGFA, gene encoding vascular endothelial growth factor-A protein; CI,
confidence interval.
\* By Chi-square test with a $2 \times 3$ contingency table.
† By Chi-square test with $2 \times 2$ contingency table.
¶ Corrected p values (Pc) were the p values corrected by multiple hypothesis tests with Bonferroni's method.

### Haplotypes constructed with the four significant SNPs in the Sherpa highlanders and non-Sherpa lowlanders

Pairwise linkage disequilibrium (LD) analysis revealed that the SNPs *r*s699947, rs833061, rs1570360, and rs2010963 were strongly in linkage in both populations (Fig 2). The haplotype comprising the Sherpa-major alleles (rs699947**C**, rs833061**T**, rs1570360**G**, and rs2010963**C**; **CTGC** haplotype) exhibited *a* significantly higher frequency in the Sherpa highlanders (0.667) than in the non-Sherpa lowlanders (0.280, P = 3.1 × 10^−5^, Fig 2, Table 3). Moreover, Sherpas carrying the **CTGC** haplotype presented a circulating VEGF-A at high altitude that was equivalent to the level in non-Sherpas with the **CTGC** haplotype at low altitude (257.62 ± 29.76 pg/mL vs. 261.39 ± 17.13 pg/mL, P = 0.91, Fig 3A). This result suggested a phenotype in which VEGF-A in Sherpa highlanders exhibits tolerance to hypoxia at high altitude. This notion was supported by the finding that the VEGF-A level in Sherpas carrying the CTGC haplotype was not significantly correlated with SpO_2_ level at high altitude (r = 0.16, P > 0.05, Fig 3B), while the VEGF-A level in non-Sherpas with the CTGC haplotype was significantly negatively correlated with SpO_2_ level at low altitude (r = −0.21, P < 0.05, Fig 3C). The other haplotypes did not show significantly different frequencies between the two groups (Table 3).

**Table 3.**
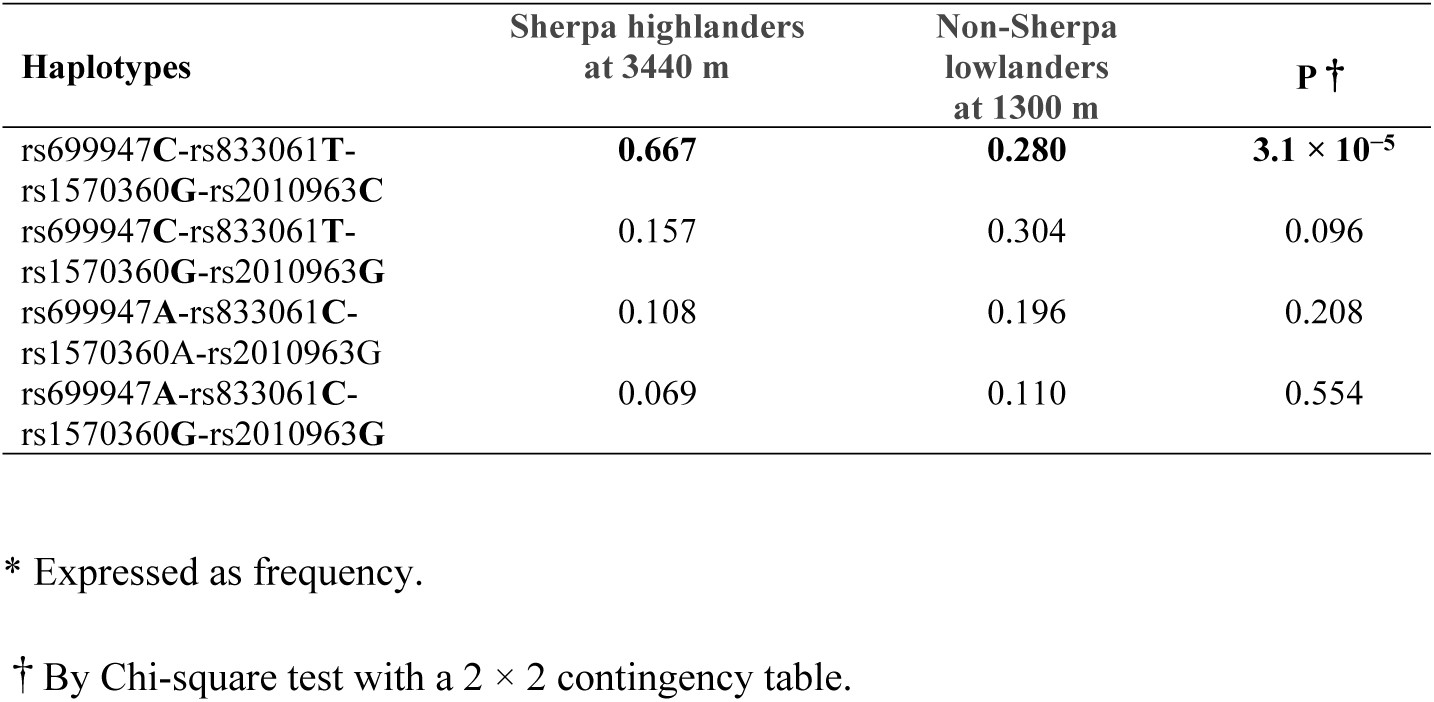
Haplotypes comprising the four significant SNPs in the Sherpa highlanders and non-Sherpa lowlanders*.

**Fig 2.**
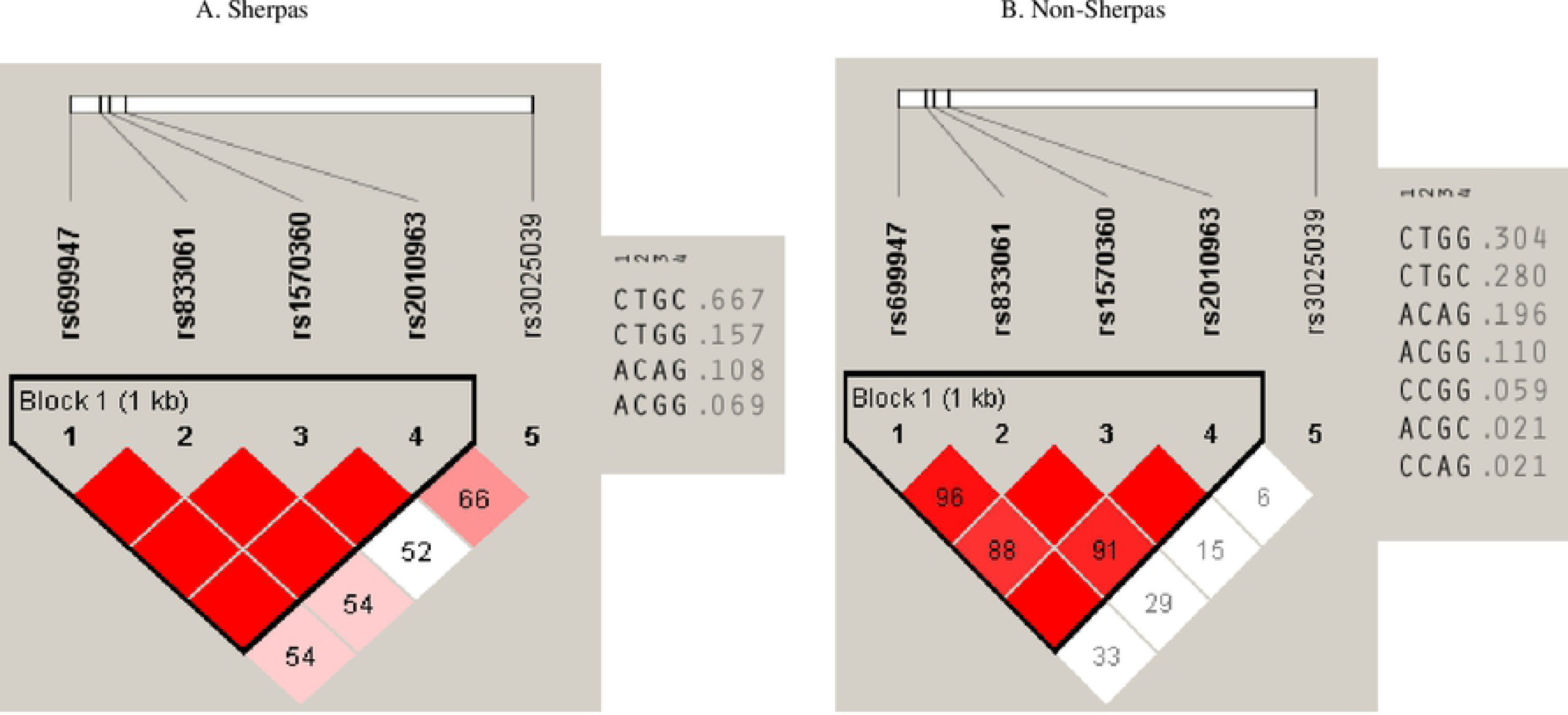
Haplotypes with the SNPs of *rs699947, rs833061, rs1570360, and rs2010963* in the Sherpa highlanders and non-Sherpa lowlanders. A) The haplotype comprising rs699947**C**, rs833061**T**, rs1570360**G**, and rs2010963**C** (**CTGC** haplotype) was prevalent among the Sherpa highlanders (frequency, 0.667). B) The frequency of the **CTGC** haplotype was 0.280 among non-Sherpa lowlanders.

**Fig 3.**
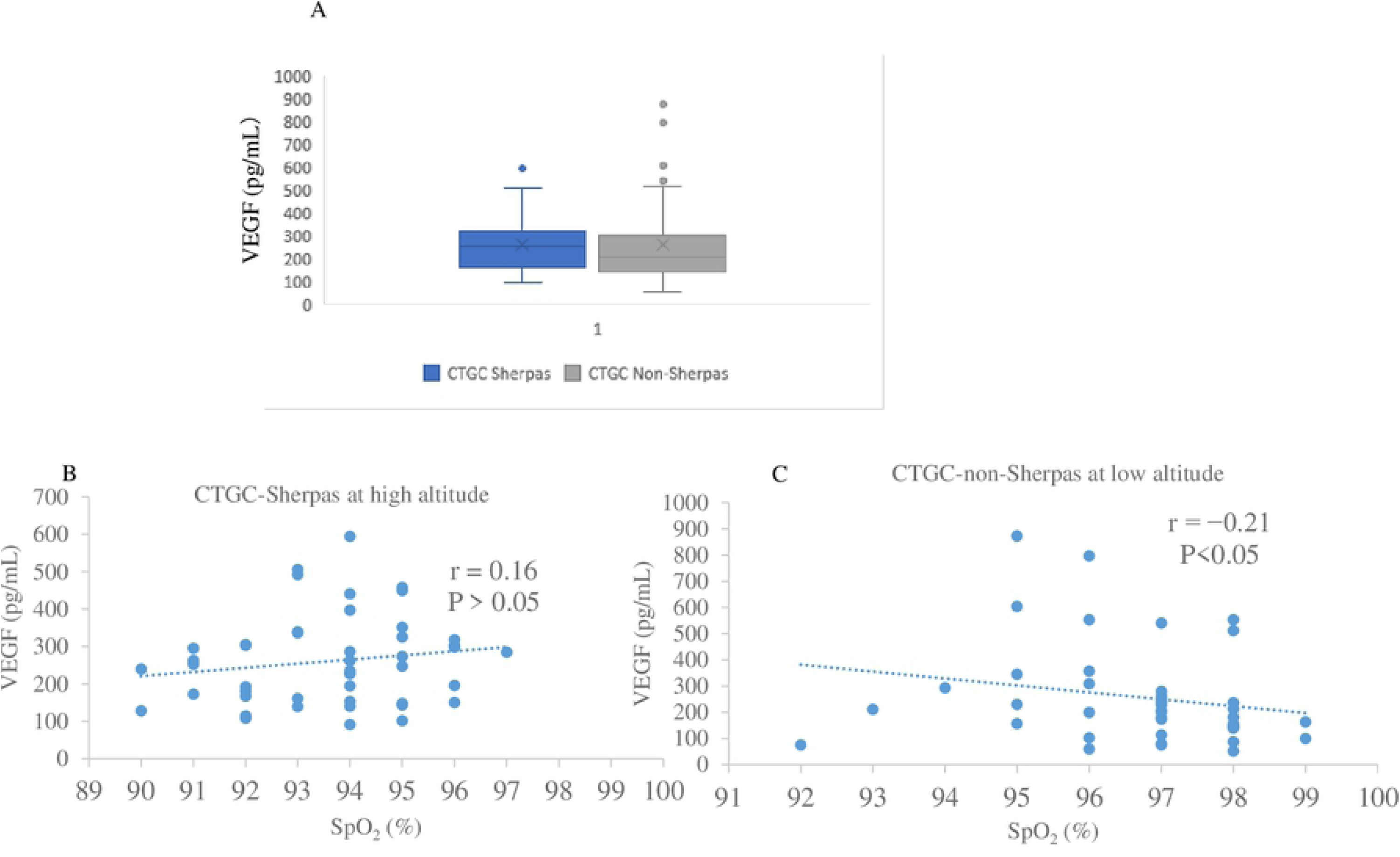
Plasma VEGF-A concentrations in Sherpa highlanders and non-Sherpa lowlanders carrying the CTGC haplotype. A) The circulating VEGF-A concentrations were similar in the two populations carrying the CTGC haplotype at different altitudes (257.62 ± 29.76 pg/mL vs. 261.39 ± 17.13 pg/mL, P = 0.91). B) VEGF-A was not significantly correlated with the SpO_2_ level in Sherpa highlanders carrying the CTGC haplotype (r = 0.16, P > 0.05). C) VEGF-A level was negatively correlated with the SpO_2_ level in non-Sherpa lowlanders carrying the CTGC haplotype (r = −0.21, P < 0.05).

### Distributions of the five examined SNPs of *VEGFA* in populations worldwide

Buroker et al. previously reported genetic information regarding the significant SNPs of *VEGFA in* Tibetan highlanders [20, 21]. Using this information, we compared the allele frequencies of these SNPs among the Sherpas, healthy Tibetans, and Tibetans with CMS (CMS-Tibetans). We found that the allele frequencies of rs699947, rs1570360, rs2010963, and rs3025039 in Sherpas were statistically consistent with those in healthy Tibetans (Table 4). However, for the SNP rs2010963, the major allele C occurred with a significantly higher frequency in Sherpas (0.667) than in CMS-Tibetans (0.521, P = 0.03, Table 4), with an odds ratio of 1.84 for the major allele C conferring strong protection against CMS in Sherpa highlanders.

**Table 4.** Allele frequencies of the SNPs rs699947, rs1570360, rs2010963, and rs3025039 in *VEGFA* among Sherpas, heathy Tibetans, and Tibetans with CMS (CMS-Tibetans)

| SNPs | Populations | Number of Subjects | Major/Minor Allele<br>number (frequency) |  | P values¶ |
| --- | --- | --- | --- | --- | --- |
| rs699947 (C/A)*<br>promoter |  |  | **C** | **A** |  |
|  | Sherpas | 51 | 84 (0.824) | 18 (0.176) | 0.94 (S vs. HT) |
|  | Heathy Tibetans | 32 | 53 (0.828) | 11 (0.172) | 0.57 (HT vs. CMST) |
|  | CMS-Tibetans | 48 | 76 (0.792) | 20 (0.208) | 0.56 (S vs. CMST) |
| rs1570360 (G/A)*<br>promoter |  |  | **G** | **A** |  |
|  | Sherpas | 51 | 91 (0.892) | 11 (0.108) | 0.76 (S vs. HT) |
|  | Heathy Tibetans | 27 | 49 (0.907) | 5 (0.093) | 0.55 (HT vs. CMST) |
|  | CMS-Tibetans | 44 | 77 (0.875) | 11 (0.125) | 0.71 (S vs. CMST) |
| rs2010963 (C/G)*<br>5'UTR |  |  | **C** | **G** |  |
|  | Sherpas | 51 | 68 (0.667) | 34 (0.333) | 0.21 (S vs. HT) |
|  | Heathy Tibetans | 30 | 34 (0.567) | 26 (0.433) | 0.58 (HT vs. CMST) |
|  | CMS-Tibetans | 47 | 49 (0.521) | 45 (0.479) | **0.03 (S vs. CMST)§** |
| rs3025039 (C/T) †<br>3'UTR |  |  | **C** | **T** |  |
|  | Sherpas | 51 | 85 (0.833) | 17 (0.167) | 0.27 (S vs. HT) |
|  | Heathy Tibetans | 29 | 52 (0.897) | 6 (0.103) | 0.85 (HT vs. CMST) |
|  | CMS-Tibetans | 32 | 58 (0.906) | 6 (0.094) | 0.18 (S vs. CMST) |

Abbreviations: S, Sherpas; HT, Heathy Tibetans; CMS, chronic mountain sickness; CMST, Tibetans with CMS.
¶ By Chi-square test with a $2 \times 2$ contingency table.
\* From Buroker J Physio Sci, 2013. (Ref. 20).
† From Buroker, Int J Hematol, 2012 (Ref. 21).
§ Odds Ratio (95% Interval Confidence) = 1.84 (1.03–3.27).

Globally, the SNPs rs699947, rs833061, and rs1570360 in the promotor and rs2010963 in the 5′-UTR of *VEGFA* exhibited *a* high degree of genetic divergence between high-altitude populations versus populations in East Asia and South Asia, as well as versus the whole global population (Fig 4). This suggested the distinct genetic variations of these SNPs in the high-altitude populations. However, rs3025039 in the 3′-UTR of *VEGFA* did not show such genetic divergence, and was thus likely to be a preserved constant SNP in populations worldwide (Fig 4).

**Fig. 4.**
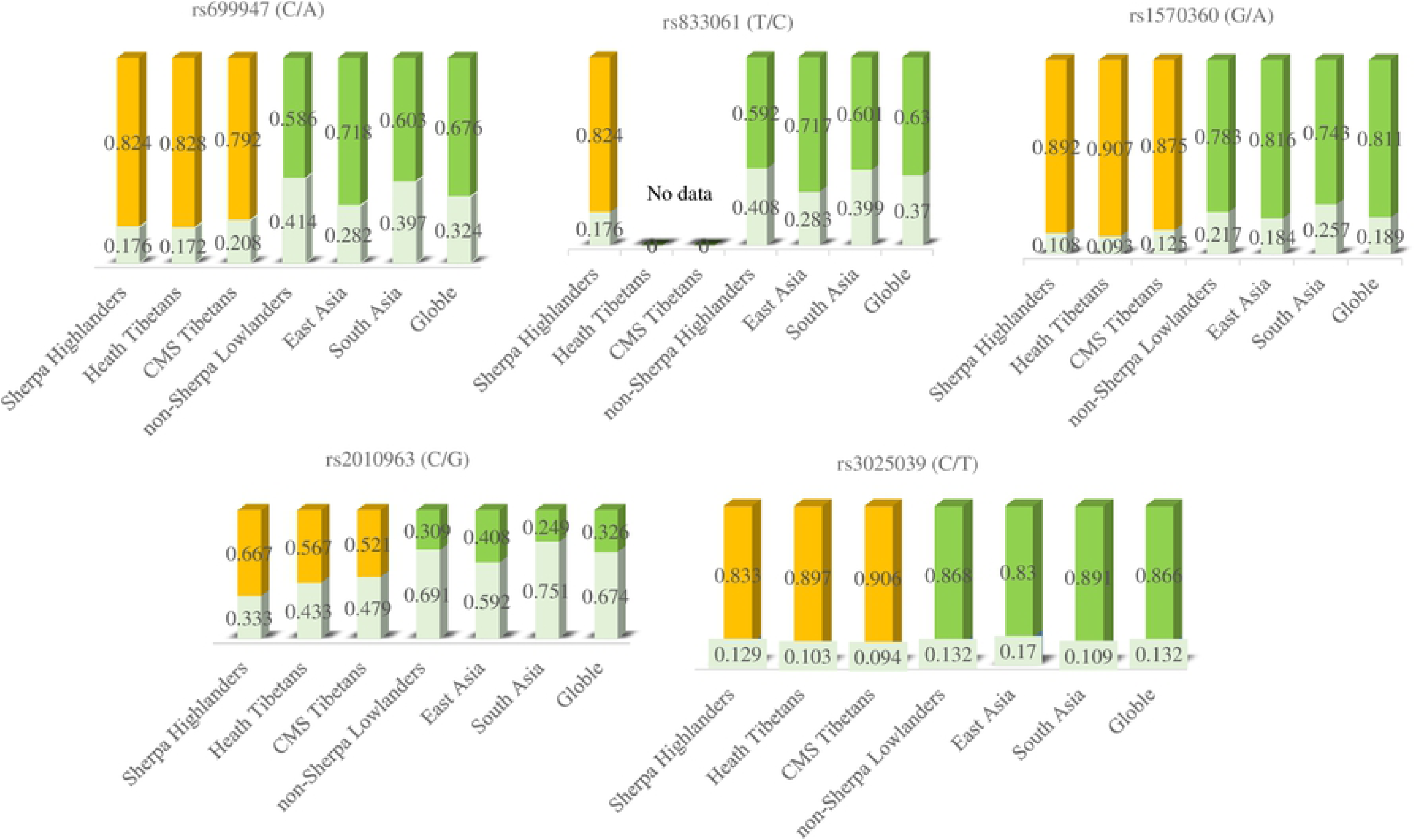
Genetic divergence of the tested SNPs in high-altitude populations versus populations in East Asia and South Asia, as well as versus the whole global population. Genetic information for healthy Tibetans and Tibetans with chronic mountain sickness (CMS) was obtained from Buroker (Ref. 20, 21). Genetic information for other populations worldwide was obtained from the 1000 Genomes Project (http://grch37.ensembl.org/Homo_sapiens/Info/Index). The populations in East Asia included Han Chinese in Beijing, China; Japanese in Tokyo, Japan; Southern Han Chinese; Chinese Dai in Xishuangbanna, China; Kinh in Ho Chi Minh City, Vietnam. The populations from South Asia included Gujarati Indian from Houston, Texas; Punjabi from Lahore, Pakistan; Bengali from Bangladesh; Sri Lankan Tamil from the UK; and Indian Telugu from the UK. Data regarding major alleles are marked with yellow for high-altitude populations and with green for low-altitude populations globally.

## Discussion

The Sherpa people are a population that exhibits adaptation to high altitude in terms of physiological tolerance to high-altitude hypoxia [4]. SpO_2_ and pulse rate are the essential physiological signs that indicate high-altitude acclimatization status. Lowlanders who ascend to a high altitude of 3500 m exhibit an average SpO_2_ of 79% and average pulse rate of 121 bpm [22]. In contrast, in our present analysis, the Sherpa highlanders presented a relatively high SpO_2_ level of 93.3 ± 0.2% and low pulse rate of 80.7 ± 1.1 bpm at a high altitude of 3440 m (Table 1). None of the Sherpa highlanders in the present study complained of CMS symptoms at high altitudes, and thus all were well adapted to a high-altitude environment. The most remarkable finding in our present study was that the Sherpa highlanders exhibited a physiological sea-level VEGF-A concentration at a high altitude. This may be a biological phenotype representing the adaptation to long-term high-altitude hypoxia in Sherpas at high altitude.

Compared to lowlanders, Sherpa highlanders exhibit a significantly higher number of capillaries per square millimeter of muscle cross-section, which would support efficient and effective diffusion of oxygen to muscles [9]. Moreover, a recent study demonstrated that Sherpas exhibit significantly higher sublingual small vessel density, capillary density, and microcirculatory flow per unit time and per unit volume of tissue compared to lowlanders at 5300 m [10]. These findings suggest the notion that peripheral vascular factors at the microcirculatory level play an important role in the process of adaptation to hypoxia in Sherpa highlanders.

VEGF-A production is mainly stimulated by hypoxia, and is upregulated in vascular smooth muscle cells and endothelial cells under hypoxic conditions [7, 23]. Studies have shown that circulating VEGF-A levels significantly increase when sea-level residents acutely ascend to high altitude [15]. Additionally, patients with CMS show significantly elevated serum VEGF-A levels [17]. Interestingly, here we found that the circulating VEGF-A level did not significantly change in response to hypoxia in Sherpas at the high altitude, and was equivalent to the level in lowlanders at the low altitude (Table 1). This suggested that VEGE-A in Sherpa highlanders was tolerant to hypoxia. This notion was further supported by the finding that VEGF-A level was not significantly correlated with the hypoxia status in Sherpas at high altitude (r = 0.13, P > 0.05, Fig 1A). In non-Sherpas at low altitude, the VEGF-A level responded to hypoxia and showed a negative correlation with hypoxia status (r = −0.27, P < 0.05, Fig 1B).

Ma et al. demonstrated that VEGF-A levels were significantly elevated in individuals susceptible to CMS at high altitudes, leading to the formation of abnormal new blood vessels in the skin and mucosa [19]. Additionally, patients with CMS showed an elevated circulating VEGF-A level with a negative correlation with SpO_2_ level [17]. Furthermore, genetic investigation revealed that the *VEGFA* expression was upregulated in patients with CMS but not in the healthy controls of high-altitude native Andeans [18]. It seems that the dysregulated overexpression of *VEGFA* induced by hypoxia may play a role in CMS pathogenesis [3], probably by the subsequent consequence of uncontrolled excessive vasculogenesis and angiogenesis that eventually interfere with the efficient oxygen diffusion in tissues [24]. We propose that VEGF-A biological pathway in Sherpa highlanders is tolerant to high-altitude hypoxia, preventing VEGF-A overproduction, and resulting in optimal physiological angiogenesis for Sherpa highlanders at high altitude. This may contribute to maintaining a sufficient capillary surface area and capillary density to restore the oxygen supply in tissues and cells under hypoxic conditions.

The Sherpa highlanders and non-Sherpa lowlanders in the present study were all Nepalese and living in Nepal. High-altitude exposure has been the major difference between these two groups, which have otherwise shared politics, economics, and cultures over the last five centuries, and have had equivalent societies and medical health services. Considering that geography is a better determinant of human genetic differentiation than ethnicity [25], we compared the genetic information regarding the rs699947, rs833061, rs1570360, rs2010963, and rs3025039 SNPs of *VEGFA* between Sherpa highlanders and non-Sherpa lowlanders. Our results revealed that the highlanders and lowlanders significantly differed in the genotype distributions and allele frequencies of rs699947 and rs833061 in the promotor region and rs2010963 in the 5′-UTR of *VEGFA* (Table 2). We additionally compared our findings with genetic data in East Asian and South Asian populations, and in the global population. The genetic divergences of SNPs in the promotor region and 5′-UTR of *VEGFA* in Sherpa highlanders were distinct among the global populations (Figure 4). Haplotype analysis also supported that the major alleles of rs699947, rs833061, rs1570360, rs2010963 in Sherpas were in genetic linkage, showing significant enrichment of the **CTGC** haplotype in Sherpa highlanders (frequency, 0.668) compared with non-Sherpa lowlanders (frequency, 0.280) (Fig 2). Sherpas carrying the CTGC haplotype exhibited a hypoxia-tolerant circulating VEGF-A phenotype (Fig 3A), and lack of correlation between VEGF-A level and SpO_2_ (Fig 3B). This **CTGC** haplotype was inherited throughout generations due to genetic linkage under the selection pressure of high-altitude hypoxic environment.

Studies of *VEGFA* SNPs in the pathogenesis of CMS have indicated that genetic variations of *VEGFA* have pathophysiological effects on CMS development [20, 26]. Here we found that the frequencies of the major alleles of rs699947, rs1570360, rs2010963, and rs3025039 in the Sherpa highlanders (0.824, 0.892, 0.667, and 0.833, respectively) were equivalent to those in heathy Tibetans dwelling on the Tibetan Plateau (0.828, 0.907, 0.567, and 0.897, respectively) (Table 4). On the other hand, the 2010963C allele frequency was significantly higher in Sherpa highlanders compared to Tibetans with CMS (0.667 vs. 0.521, P = 0.03, Table 4), suggesting that this SNP in the 5′-UTR of *VEGFA* plays a protective role against CMS in Sherpa highlanders. In Andean highlanders, the *VEGFA* expression was also significantly upregulated in patients with CMS and negatively correlated with SpO_2_ [18]. Additionally, 11 tag SNPs within a 2.2-kb region of *VEGFA* have been found to be associated with genetic variations of *VEGFA* in patients with CMS among Andean highlanders [27].

Genetic variations of *VEGFA* are apparently related to pathophysiological effects affecting CMS susceptibility, likely through targeting transcription factor binding sites (TFBS) of *VEGFA* [20, 21, 26]. Hypoxia-induced gene expression was described as a transcriptional mechanism mediated by hypoxia-responsive elements (HREs) present in the promoter region in the 5′ flanking region of *VEGFA* [28, 29]. Such HREs allow transcriptional induction of VEGF-A factor by hypoxia in several pathological states. Prior et al. reported that DNA sequence variation in the promoter region of *VEGFA* impacts the gene expression and maximal oxygen consumption in individuals before and after a standardized program of aerobic exercise training [30]. The SNPs rs699947, rs833061, rs1570360, and rs2010963 were in the TFBS in the HRE in the promotor region of *VEGFA*, with involvement in gene expression and regulation [20]. Functional analysis of the *VEGFA* 5′-UTR in hypoxic Hep3B cells has revealed that the rs699947 SNP provides a binding location for the hypoxia-induced factor (HIF)-alpha protein dimer to attach to the *VEGFA* promoter and regulate the gene [29]. The SNPs rs699947 and rs2010963 were associated with significantly altered *VEGFA* promoter activity, and responded to hypoxia exposure [13]. It seems that the genetic variations of these SNPs regulate the oxygen sensitivity of *VEGFA*, presenting a phenotype of VEGF-A hypoxia tolerance in Sherpa highlanders.

An obvious limitation of the present study was the lack of phenotypes related to capillary density and microcirculation for evaluation of angiogenesis and vasculogenesis in Sherpa highlanders at high altitude. This was due to several reasons, including the unavailability of noninvasive pathology examinations and the lack of portable equipment to carry to high altitude for fieldwork. In addition, the small sample size impeded the statistical analysis for associations of the VEGF-A levels in Sherpas at high altitude with the tested SNPs. Theoretically, the study design would have been superior if we had compared the phenotypes and genotypes between the highlanders and lowlanders at an identical height of high-altitude. Unfortunately, it was impractical to move a large number of subjects from lowland to highland, due to ethical, cultural, and logistical issues.

In summary, Sherpa highlanders at the high altitude show a physiological sea-level VEGF-A concentration, which is highly possible related to the distinct genetic background including significant variations of SNPs and haplotype patterns in the regulatory region of *VEGFA* in this population. In this setting, *VEGFA* expression is regulated in a manner exhibiting tolerance to hypoxia, promoting the production of an optimal biological level of VEGF-A at high altitudes. This enables precise angiogenesis that contributes to efficient and effective diffusion of oxygen to tissues, due to an adaptive level of capillary density and microcirculation in Sherpas at high altitude. Natural selection in a high-altitude environment for hundreds of generations would have favored *VEGFA* for adaptive advantage for Sherpa highlanders dwelling at high altitudes.

## Materials and Methods

### Ethics statement

Our study protocol was developed in accordance with the principles outlined in the Declaration of Helsinki of the World Medical Association, and was approved by the Ethics Committee of Shinshu University (Matsumoto, Japan) and the Nepal Health Research Council (Kathmandu, Nepal). The protocol was individually explained to each Sherpa highlander and non-Sherpa lowlander, and each participant gave their informed consent written in Nepalese, by signature or by fingerprint if the subject was illiterate.

### Study populations

#### Sherpa highlanders

This group comprised 51 Sherpas who lived in Namche Bazaar village (3440 m) in the Khumbu region of Nepal. The Sherpas voluntarily participated in this investigation. The Sherpa clan was identified with the Sherpa surname, and confirmed by a senior native Sherpa. All enrolled Sherpas were born and permanently resided in Namche Bazaar, and had no history of intermarriage with other ethnic groups. Information about demography, health status, altitude residence, and mountaineering history was obtained by interview. Medical interviews and physical examinations were conducted to exclude mountain sicknesses and other cardiopulmonary disorders. The SpO_2_ and pulse rate were measured using a pulse oximeter (Pulsox-3; Minolta, Osaka, Japan) with a probe connecting to a finger in Namche Bazaar village (3440 m). Venous blood samples were taken and put in tubes containing anticoagulant EDTA at high altitude (3440 m).

#### Non-Sherpa lowlanders

This group comprised 76 non-Sherpa lowlanders who lived in Kathmandu (1300 m) in Nepal. The protocol for their recruitment was the same as that followed for Sherpa highlander recruitment. In this group, SpO_2_ and pulse rate were measured, and venous blood samples were obtained at low altitude (1300 m).

### Measurement of VEGF-A concentration in plasma

Plasma VEGF-A concentration was measured using the quantitative sandwich ELISA kit (Human VEGF Quantikine; R&D Systems, Minneapolis, MN) following the manufacturer’s instructions in our laboratory at low altitude (600 m). Measurements were performed with samples in duplicate. This assay exhibited an intra-assay precision of 4.5%, and an inter-assay precision of 7.0%.

### Single-nucleotide polymorphisms

Human *VEGFA* is located on chromosome 6p21.3 and comprises eight exons that are alternatively spliced to generate isoforms of the VEGF-A protein [31]. There is considerable variation in the correlation between *VEGFA* polymorphisms and VEGF-A protein production [32, 33]. Haplotype analysis revealed that carriage of the SNPs rs833061 (in the promoter) and rs2010963 (in the 5′-UTR) significantly altered *VEGFA* promoter activity and responsiveness to biological stimuli, such as hypoxia [34]. Additionally, Prior et al. demonstrated that the VEGFA haplotype with three SNPs (rs69994 and rs1570360 in the promoter, and rs2010963 in the 5′-UTR) was associated with enhanced *VEGFA* gene expression, and with maximal oxygen consumption in individuals before and after a standardized program of aerobic exercise training [30]. Renner et al. reported that SNP rs3025039 (in the 3′-UTR of *VEGFA*) was significantly associated with altered plasma VEGF-A levels [32]. Thus, in the present study, we investigated these five SNPs: rs699947, rs833061, rs1570360 in the promoter; rs2010963 in the 5′-UTR; and rs3025039 in the 3′-UTR (Fig 5, Table 5).

**Table 5.**
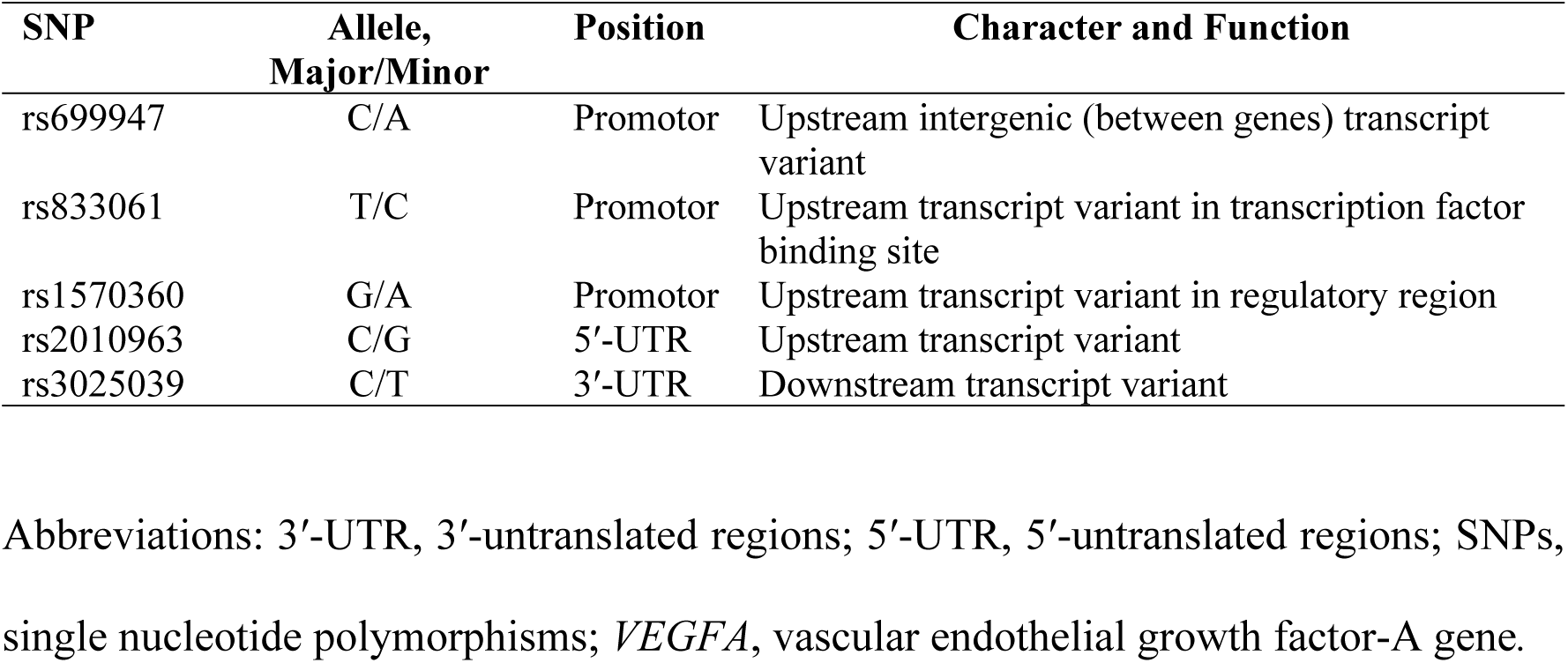
*S*ingle-nucleotide polymorphisms *of VEGFA* investigated in this study.

**Fig 5.**
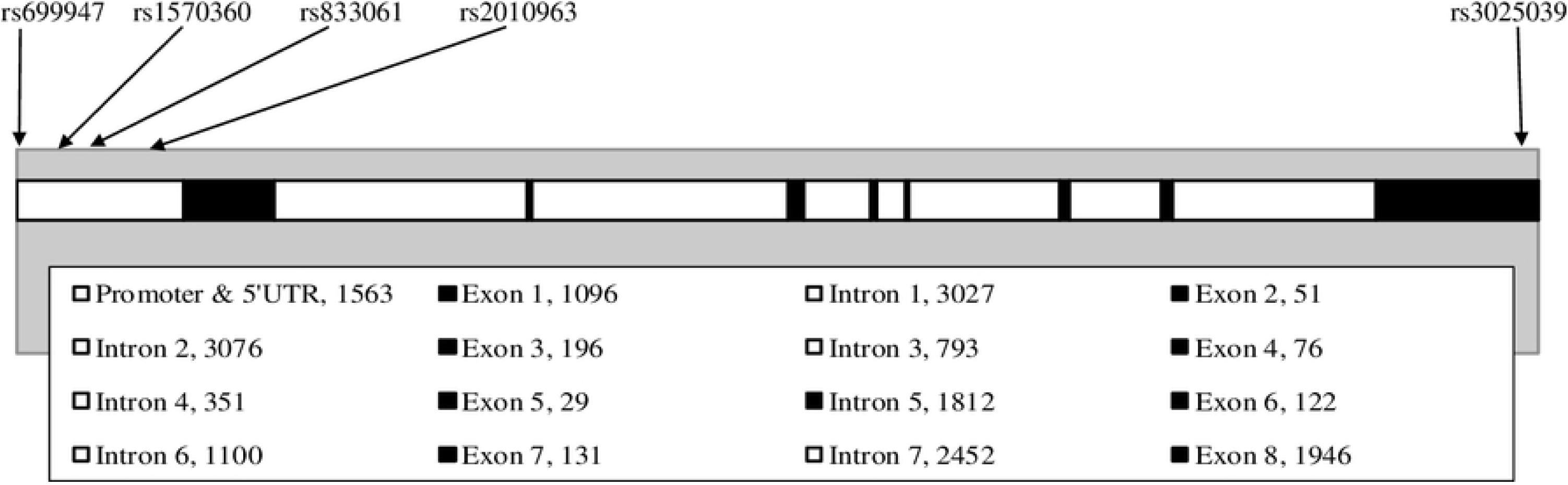
Structure of *VEGFA.* The human *VEGFA* is located on chromosome 6p21.3 and comprises eight exons. Arrows indicate the positions of the single-nucleotide polymorphisms examined in the present study. Solid boxes indicate exons, open boxes indicate introns, and numbers indicate the length in base pairs (bp).

### Genotyping

Genomic DNA was extracted from the cells in venous blood using phenol extraction in our laboratory at low altitude as previously described [5]. Allele discrimination was performed using the TaqMan® SNP Genotyping Assay with the Applied Biosystems 7500 Fast Real-time PCR System (Applied Biosystems Inc. Foster City, CA, USA) following the manufacturer’s instructions. After thermal cycling, genotype data were automatically acquired and analyzed using sequence detection software (SDS v1.3.1; Applied Biosystems, Inc.).

### Genetic data from public repository

Sherpa means “coming from East” in Sherpa’s language, referring to Tibetans from the Eastern region of Tibet Plateau, in which Sherpa origin may be traced back. Genetic evidence suggests that Tibetans are the ancestral populations of the Sherpas, whose adaptive traits for high altitude were recently inherited from their ancestors in Tibet [35]. To compare the *VEGFA* genetic variations between Sherpas and Tibetans within these high-altitude populations, we searched for genetic information regarding the five SNPs in Tibetan highlanders using English electronic databases of PubMed and Web of Science. We also obtained genetic information regarding these five SNPs in populations worldwide from the 1000 Genomes Project (http://grch37.ensembl.org/Homo_sapiens/Info/Index).

### Statistical analysis

Continuous data are expressed as mean ± standard of error (SE). To detect significant differences between the Sherpa highlanders and non-Sherpa lowlanders, we used Student’s t test for continuous data, and the Chi-square test for categorical data. The Pearson correlation was determined to assess the extent to which plasma VEGF-A level and SpO_2_ exhibited a linear relationship. The Hardy-Weinberg equilibrium was individually calculated for each SNP in the two groups using the Genepop software package [36]. We estimated the strengths of the major allele frequencies in Sherpa highlanders relative to non-Sherpa lowlanders by calculating odds ratio (OR) with the approximate 95% confidence interval (CI). P values were corrected for multiple hypothesis tests with Bonferroni’s method. The corrected P (Pc) value of less than 0.05 was considered significant.

To examine the linkage disequilibrium (LD) of the five SNPs, we used Haploview 4.2 software to derive the pairwise LD measurements and logarithm of odds of LD (D′). The haplotype was estimated according to the D′ values, based on the maximum likelihood to generate strong LD of the SNPs [37].

## Acknowledgments

We are grateful to all the Sherpa highlanders and non-Sherpas lowlanders for their kind participation in this study. We thank Doctors of Amit Arjyal, Pritam Neupane, Anil Pandit, and Dependra Sharma for their assistances during sample collections in the Sherpa village and Kathmandu in Nepal. We appreciated the cooperation from the Nepal Health Research Council (Katmandu, Nepal).

